# BioModelsML: Building a FAIR and reproducible collection of machine learning models in life sciences and medicine for easy reuse

**DOI:** 10.1101/2023.05.22.540599

**Authors:** Divyang Deep Tiwari, Nils Hoffmann, Kieran Didi, Sumukh Deshpande, Sucheta Ghosh, Tung V. N. Nguyen, Karthik Raman, Henning Hermjakob, Rahuman Sheriff

**Affiliations:** EMBL-EBI, Wellcome Genome Campus, Hinxton, Cambridgeshire, CB10 1SD, UK; Forschungszentrum Jülich GmbH, Germany; Cardiff University, UK; Heidelberg Institute for Theoretical Studies-HITS gGmbH, Germany; Indian Institute of Technology Madras, Chennai, India

## Abstract

Machine learning (ML) models are widely used in life sciences and medicine; however, they are scattered across various platforms and there are several challenges that hinder their accessibility, reproducibility and reuse. In this manuscript, we present the formalisation and pilot implementation of community protocol to enable FAIReR (Findable, Accessible, Interoperable, Reusable, and Reproducible) sharing of ML models. The protocol consists of eight steps, including sharing model training code, dataset information, reproduced figures, model evaluation metrics, trained models, Dockerfiles, model metadata, and FAIR dissemination. Applying these measures we aim to build and share a comprehensive public collection of FAIR ML models in the BioModels repository through incentivized community curation. In a pilot implementation, we curated diverse ML models to demonstrate the feasibility of our approach and we discussed the current challenges. Building a FAIReR collection of ML models will directly enhance the reproducibility and reusability of ML models, minimising the effort needed to reimplement models, maximising the impact on the application and significantly accelerating the advancement in the field of life science and medicine.

## Introduction

Machine Learning (ML) models have become widespread in life sciences and medical research. However, the scattered nature of these models across various platforms like personal websites, GitHub, bitbucket, and supplementary material, makes it challenging to find, access and reuse them. In addition, the published results of ML models are often difficult to reproduce due to various factors including missing metadata, training data, required libraries, or the code used. A survey showed that about 70% of scientists across all major branches of science could not reproduce others’ results, and 50% could not reproduce their own results, highlighting the reproducibility crisis in science (Baker, 2016). Furthermore, our recent study shows that computational biology models are not immune to this reproducibility crisis, with about half of the published models from various domains in life science being affected (Tiwari *et al*., 2021).

Thus, addressing the challenges in public sharing and reproducibility of ML models is essential. ML models should faithfully reproduce their predictions when all relevant information is provided. A study that evaluated 511 scientific papers in various subfields of machine learning indicated that compared to other areas, machine learning for healthcare falls behind in reproducibility metrics like the accessibility of datasets and codes (McDermott *et al*., 2021). Data leakage is considered an important factor to contribute to the reproducibility crisis in machine learning [4]. Data leakage refers to a situation where a model is exposed to information from evaluation data during the training process, resulting in an overestimation of accuracy (Gibney, 2022). Hence, the sharing of code, data, and random seeds used in the training of an ML model is imperative to allow third parties to evaluate and achieve technical reproducibility (Heil *et al*., 2021).

DOME, a community guideline for reporting supervised machine learning–based analyses applied to biological studies, has been developed by members and collaborators of the ELIXIR Machine Learning Focus Group. It includes a set of recommendations for reporting details on the choice of methods and parameters for Data, Optimization, Model, and Evaluation (Walsh *et al*., 2021), which mainly address the aspects of reproducibility of machine learning models. The Kipoi database was established to allow the exchange of machine learning models in genomic research to facilitate reproducible research (Avsec *et al*., 2019). Developing comparable repositories for other areas of life science and medical research would be a difficult task.

Building upon the existing efforts to promote reproducible ML model reporting and exchange, FAIR data sharing principles (Wilkinson *et al*., 2016), and the concept of centralised curation, we developed a community-based protocol to share ML models in a reproducible and reusable manner. In this manuscript, we present these protocols together with use cases and infrastructure to foster efficient adoption and implementation. Our protocols and workflows will enable the building and sharing of findable, accessible, interoperable, reusable (FAIR) and reproducible ML models.

Applying these principles, our long-term goal is to build a large public collection of manually-verified, free and open-source FAIR ML models by extending the BioModels repository (Malik-Sheriff *et al*., 2020). BioModels, an ELIXIR deposition database of mechanistic models of biological and medical systems, is hosted at EMBL-EBI and accessed by about 51,000 unique users (IPs) annually. With the proposed extension, BioModels will cater to the growing need for reproducible and reusable ML models in the scientific community.

In essence, our approach will reduce the effort required to reimplement and maximise the impact of ML models benefiting both model builders and users in the field of life science and medicine ultimately leading to enhanced research and application outcomes.

### Eight steps to make ML models FAIReR

An important goal of the ML community is to make ML models FAIR (Wilkinson *et al*., 2016) and reproducible (FAIReR). When details on the data, code and conditions are shared, an ML model should faithfully reproduce the predictions using the same input datasets; however, this is not often assessed during the peer-review process (Kapoor and Narayanan, 2022). Full reproduction of an ML model can take from weeks to months and a peer reviewer may not have time to fully scrutinise it (Gibney, 2022). Building upon the existing BioModels curation procedures (Malik-Sheriff *et al*., 2020), we propose a concept where the model author and independent community curators can collaboratively work together to enable FAIReR sharing of ML models in eight steps. An ML modeller is recommended to perform all steps except steps 3 and 4, which requires a curator’s involvement. However, a curator from the community can perform all eight steps independently to make the model FAIReR and to gain recognition for their curation work.

Table 1 summarises a checklist with tasks, the corresponding FAIReR metrics and the primary person responsible (ML modeller or curator). We propose a centralised community reproducibility assessment and sharing of ML models through a public repository such as BioModels. This approach will provide a pragmatic and scalable solution to improve the accessibility of code, data and conditions and a mechanism for improving the discoverability, interoperability, reproducibility and reusability of the ML models.

**Table 1:** Checklist to make ML models FAIR and reproducible.

| Criteria | Findability | Accessibility | Interoperability | Reusability | Reproducibility | Primary responsibility (Author and Curator) |
| --- | --- | --- | --- | --- | --- | --- |
| **Training Code:** All relevant code files to train ML model. |  | A |  | R | eR | Au |
| **Dataset:** Link to datasets used for model training and validation. | F | A |  | R | eR | Au |
| **Reproduced Figure:** Figures that are successfully reproduced. |  |  |  |  | eR | Cu |
| **Evaluation:** Table of model evaluation metrics (Accuracy, Sensitivity, Specificity, Precision etc). |  |  |  | R | eR | AuCu |
| **Trained Model:** Trained model in ONNX format. |  |  | I | R | eR | Au |
| **Docker for Building Model:** Dockerfile (with ReadMe file) - to train and build the model |  |  |  | R | eR | Au |
| **Docker for applying trained Model:** Dockerfile to load the trained ONNX model and run the prediction / inference. |  |  | I | R | eR | Au |
| **Model Metadata:** annotation file ('xml' or 'csv') with RDF containing relevant Ontology terms for methods (EDAM) and other biological features (GO, NCIT, UBERON, CellOntology, etc). | F |  | I | R |  | Au |
| **Dissemination:** Submission of model codes and trained models to BioModels repository. | F | A |  | R |  | Au |

#### Step 1: Sharing model training code

To ensure the reproducibility of ML models, several key elements need to be included. Firstly, all relevant code files used to train the ML model should be made available, including scripts to pre-process data, train the model, and evaluate its performance. This should also include the random seed used for training when relevant. This code should be well-documented and easily understandable, with comments explaining each step of the process.

#### Step 2: Sharing dataset information

In addition to the training code, a link to the dataset used for model training and validation should also be provided. The dataset should be openly accessible and adequately labelled, with information on how it was obtained and any pre-processing steps taken. Any new data should be appropriately shared following FAIR principles, in a relevant data repository. This will enable other researchers to reproduce the model and evaluate its performance on the same data.

#### Step 3: Reproducing model predictions

The model predictions should be independently reproduced by the curator. To demonstrate the reproducibility of the model, figures from the original reference manuscript that are successfully reproduced should also be publicly shared. These figures should be accompanied by a description of how they were generated, including any code used to create them. This will enable other researchers to verify that the model produces the expected results.

#### Step 4: Sharing model evaluation metrics

To evaluate the performance of the model, a table of model evaluation metrics should be provided. This table should include common metrics such as accuracy, sensitivity, specificity, and precision. The calculation of these metrics should be clearly documented.

#### Step 5: Sharing trained model

The trained model should be made available in either native format or ONNX (Open Neural Network Exchange) format when possible. ONNX is a widely used open-source format designed to foster interoperability between different machine learning frameworks, and therefore, will allow users to interchange ML models between various machine learning frameworks and tools (Bai, Fang and Ke, 2019). This format should be accompanied by clear instructions on how to load and run the model to make predictions.

#### Step 6: Sharing Dockerfiles

To simplify the process of reproducing and reusing ML models, Dockerfiles should also be provided. These files should include a ReadMe file with instructions on how to use them, as well as any necessary dependencies. The first Dockerfile should allow the user to generate a Docker image to train and build the model with both original data from the modeller as well as new data from the users. The second Dockerfile should allow the user to generate a Docker container to load the trained model from Step 5 and calculate predictions using the original datasets to test reproducibility as well as to apply the model to new datasets to create new predictions. The Dockerfile provides an opportunity to generate a standardised environment for running the model, ensuring that the models can be reproduced and applied to different datasets. When Docker images are available in a public repository, they should be appropriately cross-referenced.

#### Step 7: Sharing model metadata

To ensure the FAIR sharing of machine learning models, one of the key requirements is the provision of model metadata. Model annotation in RDF (Le Novère *et al*., 2005) containing cross-references to controlled vocabularies, including relevant ontology terms for methods from EDAM (Ison *et al*., 2013; Black *et al*., 2021), as well as for other biological features from GO (Ashburner *et al*., 2000), NCIT (Golbeck *et al*., 2003), UBERON (Mungall *et al*., 2012), and CellOntology (Diehl *et al*., 2016), and standard resources, tools and datasets is critical. Currently, RDF triples containing annotation files can be uploaded as XML or CSV files to BioModels. The metadata will enable other researchers and modellers to find and identify the model based on its specific characteristics, such as the type of ML algorithm used or the biological system it was trained on, tools used, input data types, predicted outputs and other recommendations from DOME guidelines.

#### Step 8: FAIR dissemination

Dissemination of the model is also crucial for making it findable and accessible. One effective way to disseminate the model is to submit the model code, the trained model, cross-reference to original code, datasets, reproduced figure(s), evaluation metrics, Dockerfiles for training and (pre-) trained models along with model metadata to the BioModels repository. BioModels can index machine-readable annotations of the model and therefore significantly improves its discoverability through various search engines, including BioModels itself, EBI search (Madeira *et al*., 2022), OmicsDI (Perez-Riverol *et al*., 2017), and Google datasets. Furthermore, models hosted in BioModels can be accessed globally either via the web interface or programmatically, making them highly accessible to other researchers.

By following these steps and fulfilling all these elements in the Table 1, the reproducibility and the reusability of the model can be ensured, allowing other researchers to validate its results and apply the model for new predictions.

### Incentivized Community Curation

FAIReR sharing of ML models to create a comprehensive public collection of ML models in life science and medicine is a formidable task that cannot be accomplished by any single group or organisation. To address this challenge, we propose a community curation approach to invite ML modellers to contribute to the collection. The community curation effort can be harmonised by providing modellers with detailed protocols and workflows (Figure 1) for standardised model reproduction and sharing. Through this approach, we aim to facilitate the dissemination of published models through BioModels and make them freely available to the broader scientific community. To publicly acknowledge the contribution of the community contributors, we will integrate the BioModels platform with ORCID and APICURON (Hatos *et al*., 2021) to allow the modellers and curators to claim credit for their contribution, similar to claiming a publication. To guarantee that the shared ML models comply with the recommended guidelines, BioModels would carry out quality control measures before public release and provide metrics to reflect the compliance and quality of curation.

**Figure 1:**
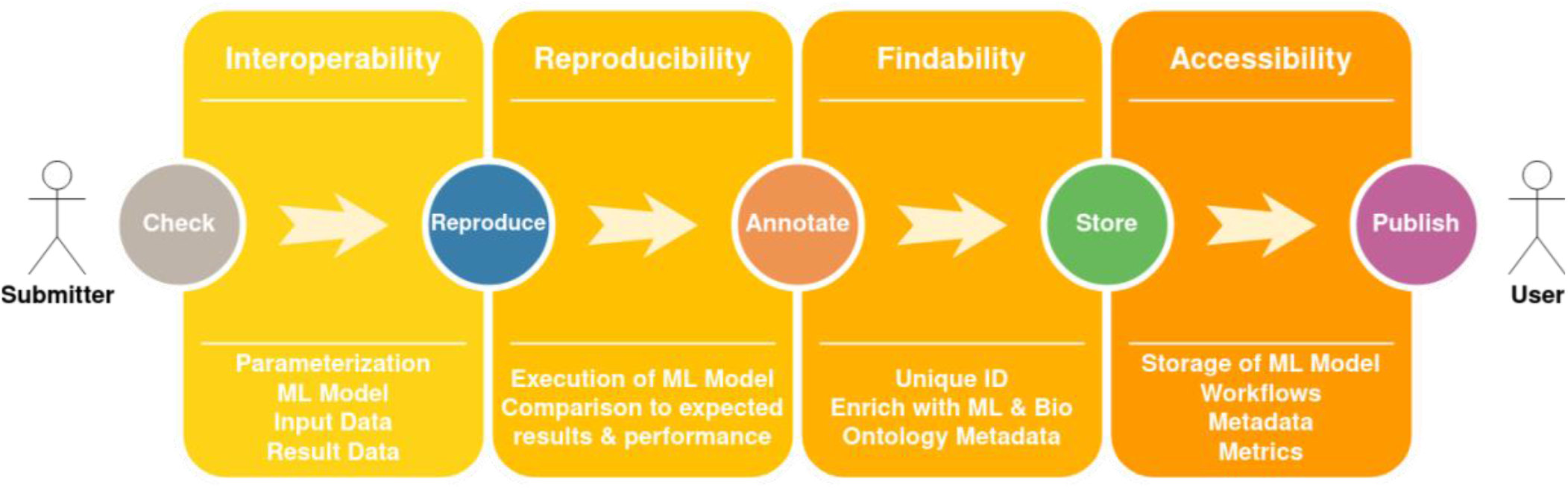
BioModels submission and curation workflow, mapped to the FAIR principles. Machine learning models can be included either in their native format or in ONNX format. Alternatively, steps to build/train native models can be included within the provided Dockerfiles.

### Pilot Implementation

Using the protocol developed above, we initiated our effort to build a FAIReR collection of ML models in BioModels, through a pilot implementation of published ML models. We developed a workflow (Figure 1) to curate models from published literature and demonstrate the feasibility of the approach. We curated six models (Table 1) from diverse domains, ranging from prediction of protein structure to coding sequence to response of patients to therapy, using a variety of methodologies from deep learning to random forests to NLP approaches, and built on tools such as TensorFlow, PyTorch and sci-kit learn. We will continue to expand this collection and all the models can be accessed through a link to the tag [‘FAIR-AIML’ in BioModels](https://www.ebi.ac.uk/biomodels/search?query=submitter_keywords:FAIR-AIML&domain=biomodels).

Reproducing and sharing FAIR ML models can take from weeks to months depending on the complexity of the model, missing information, technical challenges and time taken to communicate with the authors. Table 2 provides evidence that the recommendations outlined in Table 1 are feasible in practice, with only a few exceptions. Continuous efforts are being made to resolve these exceptions and show the recommendations fully achievable in the future. In many cases, reproducing the model initially posed various challenges, including outdated scripts and missing dependencies. Nevertheless, after consulting with the authors, these issues were successfully addressed, where necessary, allowing reproduction of all models as described in the manuscript (see Supplementary Table 1 for details). By verifying and reproducing the models, curators shared all working files through BioModels, enabling easy reuse by the public users. This approach eliminates the need for modellers to respond to numerous requests for support, leading to a more streamlined and efficient process.

**Table 2:**
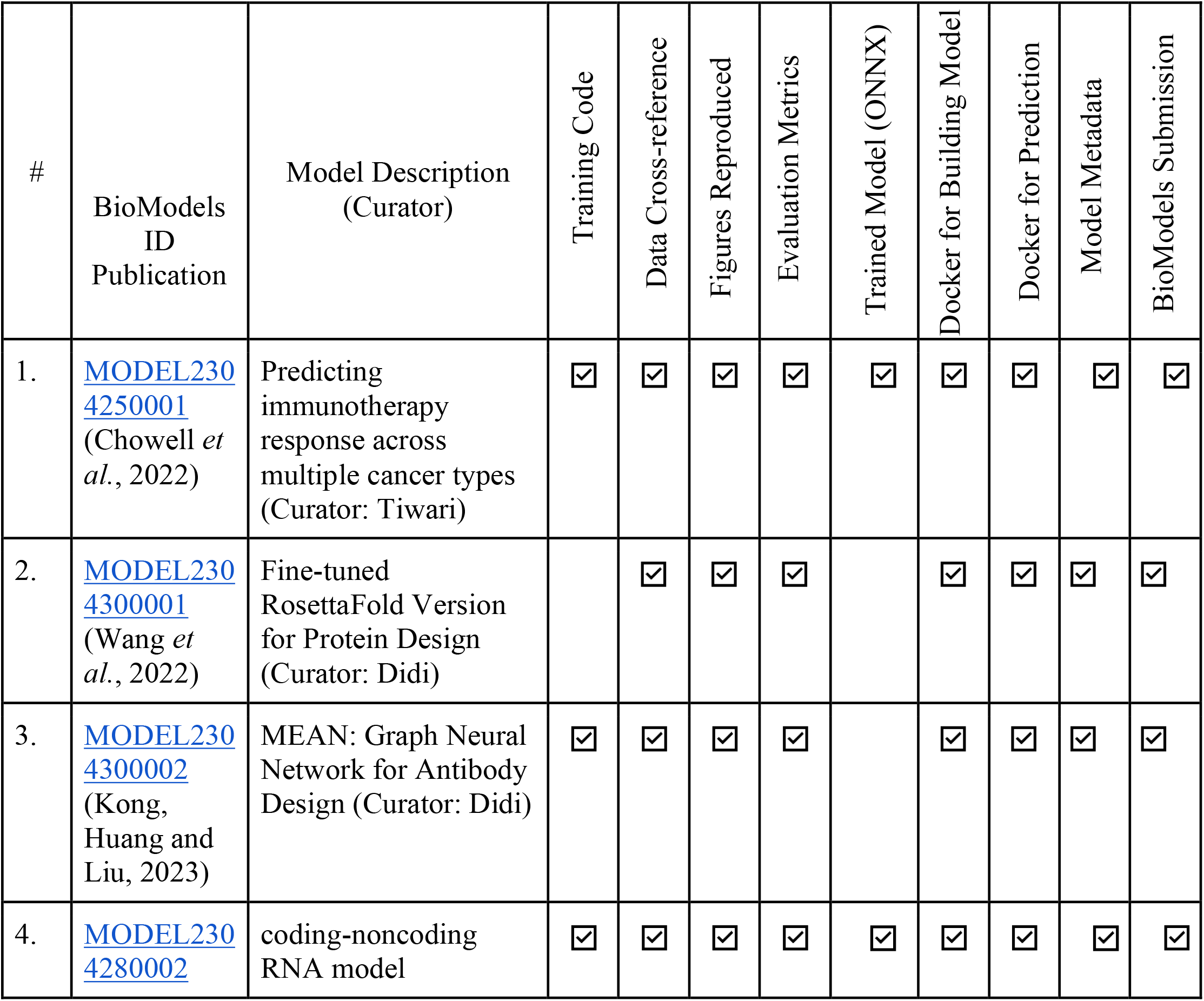
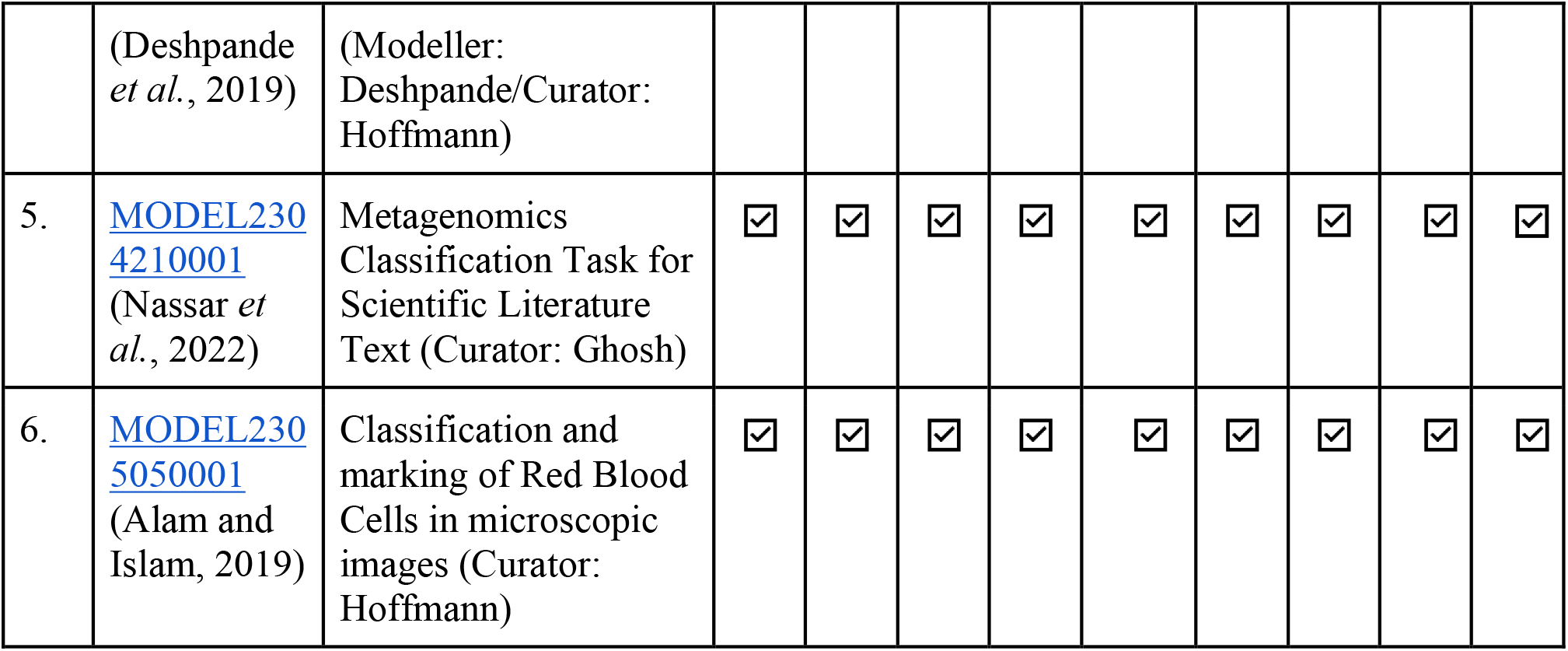
A list of use case FAIReR machine learning models in BioModels.

| # | BioModels ID Publication | Model Description (Curator) | Training Code | Data Cross-reference | Figures Reproduced | Evaluation Metrics | Trained Model (ONNX) | Docker for Building Model | Docker for Prediction | Model Metadata | BioModels Submission |
| --- | --- | --- | --- | --- | --- | --- | --- | --- | --- | --- | --- |
| 1. | MODEL2304250001<br>(Chowell *et al.*, 2022) | Predicting immunotherapy response across multiple cancer types (Curator: Tiwari) | ☒ | ☒ | ☒ | ☒ | ☒ | ☒ | ☒ | ☒ | ☒ |
| 2. | MODEL2304300001<br>(Wang *et al.*, 2022) | Fine-tuned RosettaFold Version for Protein Design (Curator: Didi) |  | ☒ | ☒ | ☒ |  | ☒ | ☒ | ☒ | ☒ |
| 3. | MODEL2304300002<br>(Kong, Huang and Liu, 2023) | MEAN: Graph Neural Network for Antibody Design (Curator: Didi) | ☒ | ☒ | ☒ | ☒ |  | ☒ | ☒ | ☒ | ☒ |
| 4. | MODEL2304280002 | coding-noncoding RNA model | ☒ | ☒ | ☒ | ☒ | ☒ | ☒ | ☒ | ☒ | ☒ |
|  | (Deshpande *et al.*, 2019) | (Modeller: Deshpande/Curator: Hoffmann) |  |  |  |  |  |  |  |  |  |
| 5. | MODEL2304210001<br>(Nassar *et al.*, 2022) | Metagenomics Classification Task for Scientific Literature Text (Curator: Ghosh) | ☑ | ☑ | ☑ | ☑ | ☑ | ☑ | ☑ | ☑ | ☑ |
| 6. | MODEL2305050001<br>(Alam and Islam, 2019) | Classification and marking of Red Blood Cells in microscopic images (Curator: Hoffmann) | ☑ | ☑ | ☑ | ☑ | ☑ | ☑ | ☑ | ☑ | ☑ |

## Discussion

This manuscript presents the protocol developed to promote FAIReR sharing of ML models in life sciences and medicine, along with our pilot implementation. With the protocol developed after reviewing various studies, we initiated our pilot effort to build a FAIReR collection of ML models in the BioModels database. In this study, we demonstrate the feasibility of our approach by curating six peer-reviewed published ML models in the BioModels database (Table 2).

Although our approach is practically achievable and scalable, there are still some minor issues that need to be addressed. One such issue is the conversion of models to ONNX, which remains challenging in some cases. It is generally easier to generate an ONNX model immediately within the respective ML framework used by the authors, while post hoc conversion may prove difficult in some cases. These issues could be solved by providing clear instructions on which formats, data, and information are required when the ML framework native formats are exported to maximise compatibility with ONNX.

Other potential obstacles include the necessary custom conversion of input and output data, e.g., to pre-process images for annotation and to post-process and map classified regions back onto the input images. Also, for more complex models, the ONNX-based predictions may sometimes deviate from the ones generated with the native model. This was the case for the YoloV2-based blood cell counting dataset, but only led to minor differences in classification performance. At the moment, it is not clear if those differences may be attributed to implementation differences for certain operators and numerical precision supported by the respective ONNX runtime implementation. ONNX is sometimes not suitable for more complex use cases for example Wang2022, and Kong 2023. ONNX is often behind the current state of the ML landscape making it hard to implement newly published models with it at the start. For this reason, the newer parts in the torch library used in the Kong 2023 cannot be converted into ONNX format.

Another challenge is the potential size of ONNX models, which can be over a gigabyte (GB) in some cases. Therefore, the BioModels platform is currently being enhanced to support efficient hosting and dissemination of such models. Additionally, ML model reproduction is a time-consuming and laborious process, and community curation can be adapted by bringing in experts from specialised domains. To facilitate this, the BioModels platform is being enhanced to enable ORCID and APICURON-based acknowledgement of modellers’ and curators’ contributions. A star-based system to report the level of curation and level of compliance with the DOME recommendations and FAIReR ML checklist is also planned to be implemented in BioModels soon.

Dockerfiles specifically for (a) training a model or (b) executing a pre-trained native or ONNX model are shared, to facilitate reproducible building and application of the ML models. In our approach, only Dockerfiles are shared, not container images, as this strategy is essential to optimise data storage in BioModels. Any pre-built Docker images available through services such as Hugging Face can be linked through the BioModels’ model description page. It is also crucial to link the original models when they are publicly available through repositories such as GitHub or Bitbucket.

One of the key aspects of our infrastructure is to support version-controlled community contributions of FAIReR models, allowing the contributor to provide independent verification of the ML models and gain recognition for their effort through the ORCID and APICURON platforms. We invite community members to join this effort and curate models following the guidelines and use cases presented in the manuscript.

An open-source and free collection of (re)usable ML models will greatly advance life science and clinical research and application. Researchers or clinicians can search and retrieve all models of their interest and filter them to identify a suitable model based on input and output types, performance metrics, and clinical conditions of the cohort. For example, a clinician aiming to identify a tumour in mammograms should be able to search relevant keywords such as breast cancer and filter relevant models that have input type as an image and output type as disease grade. The clinician can further select verified ML models with high-performing evaluation metrics. We believe that our approach will pave the way for the efficient reuse of ML models and greatly enhance their potential for gainful application in the field of life sciences and medicine.

## Acknowledgements

DDT, TN, HH, RMS acknowledge EMBL core funding.

NH, SG, SD, RMS acknowledge support from ELIXIR BioHackthon 2022.

DDT acknowledges the post-baccalaureate fellowship from RBCDSAI. KR acknowledges funding from RBCDSAI.

