## Supplementary Material for "BioModelsML: Building a FAIR and reproducible collection of machine learning models in life sciences and medicine for easy reuse"

*22 May 2023*

*Divyang Deep Tiwari[1,5], Nils Hoffmann[2], Kieran Didi[1], Sumukh Deshpande[3], Sucheta Ghosh[4], Tung V. N. Nguyen[1], Karthik Raman[5], Henning Hermjakob[1,6], Rahuman Sheriff [1,6]*
**

*[1]EMBL-EBI, Wellcome Genome Campus, Hinxton, Cambridgeshire, CB10 1SD, UK*
*[2]Forschungszentrum Jülich GmbH, Germany*
*[3]Cardiff University, UK*
*[4]Heidelberg Institute for Theoretical Studies-HITS gGmbH, Germany*
*[5]Indian Institute of Technology Madras, Chennai, India*
*[6] Corresponding authors*

*Supplementary Table 1: Overview of use cases and corresponding BioModels submissions, detailing the algorithms used, the approximate effort for reproduction of their results, the necessity of adaptations of original code, the input data type for each model and whether authors needed to be contacted. [a] Original reproduction of the native model took 1 day, 9 days for the ONNX model and implementation of compatible pre- and post-processing.*

| Use Case # | BioModels ID Publication | Model Algorithm | Machine/Deep Learning Library Used | Time Taken to Reproduce (Working Days) | Original Code Modified | Reproduced Metrics | Data Type (Tabular, Text, Image) | Authors Contacted |
|---|---|---|---|---|---|---|---|---|
| 1. | MODEL2304250001 (Chowell *et al.*, 2022) | Random Forest | Scikit-Learn | 20 | Yes | Sensitivity, Specificity, Accuracy, PPV, NPV | Tabular | Yes |

| | | | | | | | | |
|---|---|---|---|---|---|---|---|---|
| 2. | MODEL2304300001 (Wang *et al.*, 2022) | Neural network, including SE(3)-equivariant transformer | PyTorch | 10 | Yes | RMSD, pLDDT | Protein Structure Data (PDB Files) | Yes |
| 3. | MODEL2304300002 (Kong, Huang and Liu, 2023) | Graph Neural Network | PyTorch | 5 | Yes | RMS, AAR (amino acid recovery) | Protein Structure Data (PDB Files) | Yes |
| 4. | MODEL2304280002 (Deshpande *et al.*, 2019) | Random Forest | Scikit-Learn | 3 | Yes | Precision Recall F1-Score Support | Text | Yes |
| 5. | MODEL2304210001 (Nassar *et al.*, 2022) | DeepLearning & Random Forest | TensorFlow 2, Scikit-Learn | 3 | Yes | Prediction Probability | Text | Yes |
| 6. | MODEL2305050001 (Alam and Islam, 2019) | DeepLearning, CNN, Darkflow, Tiny YoloV2 | TensorFlow 2 | 10[a] | Yes | Annotation area centres Total numbers of annotated cells | Image (Microscopy) | No |

### Use Case #1: Chowell 2022, Random Forest model to predict efficacy of immune checkpoint blockade across multiple cancer patient cohorts

In this work, we reproduced a machine learning model that has been developed to predict the response to Immune Checkpoint Blockade (ICB) treatment across 16 different cancer types. The dataset was obtained from the MSK-IMPACT cohort and contains various biological information for 1479 patients treated with ICB. The model also provides a quantitative estimation of the most important features for the predictions. The cancer types have been classified into three major categories: melanoma, NSCLC (Non-Small Cell Lung Cancer), and Others (the remaining 14 categories).

Upon downloading the dataset, it was discovered that the column names in the data do not match with the column names in the original code. Therefore, necessary adjustments have been made to the original code to align it with the dataset. A Random Forest algorithm was trained on a set of 16 features (referred to as RF16) using the Scikit-Learn library for the Python programming language. The training and test prediction results were successfully reproduced.

Additionally, three figures (feature contribution plot, ROC plots for the training and test datasets), and a confusion matrix were successfully reproduced and included in the submission. Subsequently, the Scikit-Learn model was exported in the ONNX format, and a Docker file was created for training the model, as well as another Docker file for using the ONNX model for predictions. Finally, a model metadata file was developed in CSV format, and the submission was made to the BioModels repository (see Supplementary Fig.1-4).

The model files can be downloaded from the BioModels repository with model identifier BIOMD0000001066. To make predictions using your own data, you can directly execute the Dockerfile located in the "dockerICB_predict" folder. This Dockerfile is designed to enable users to make predictions using their specific data. If you wish to retrain the model, you can do so by executing the Dockerfile in the "dockerICB_train" folder. This Dockerfile is specifically created to facilitate model training. By following these steps, users can either make predictions with their own data or retrain the model according to their requirements.

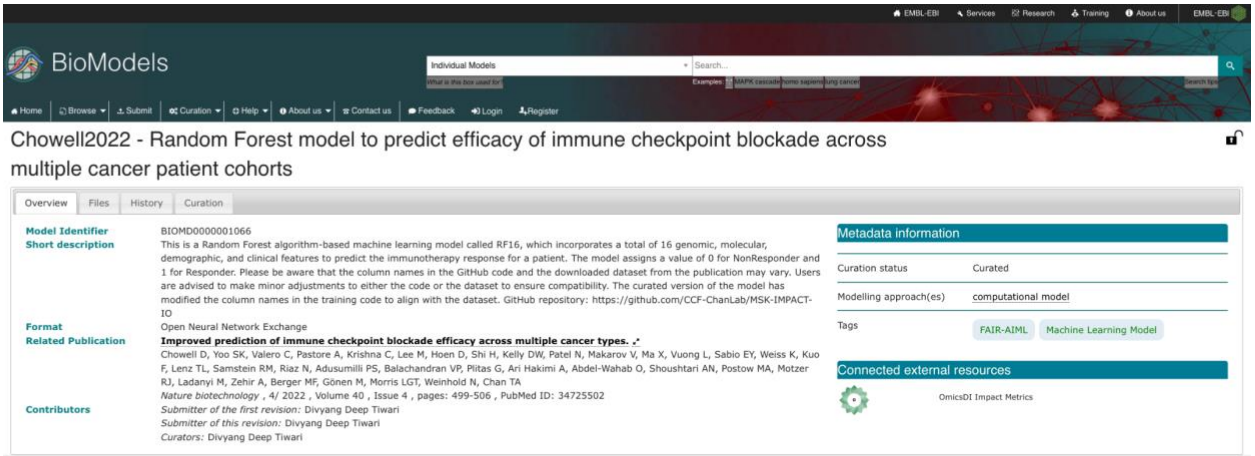


*Supplementary Figure 1: Overview Tab of Chowell2022 submitted model.*

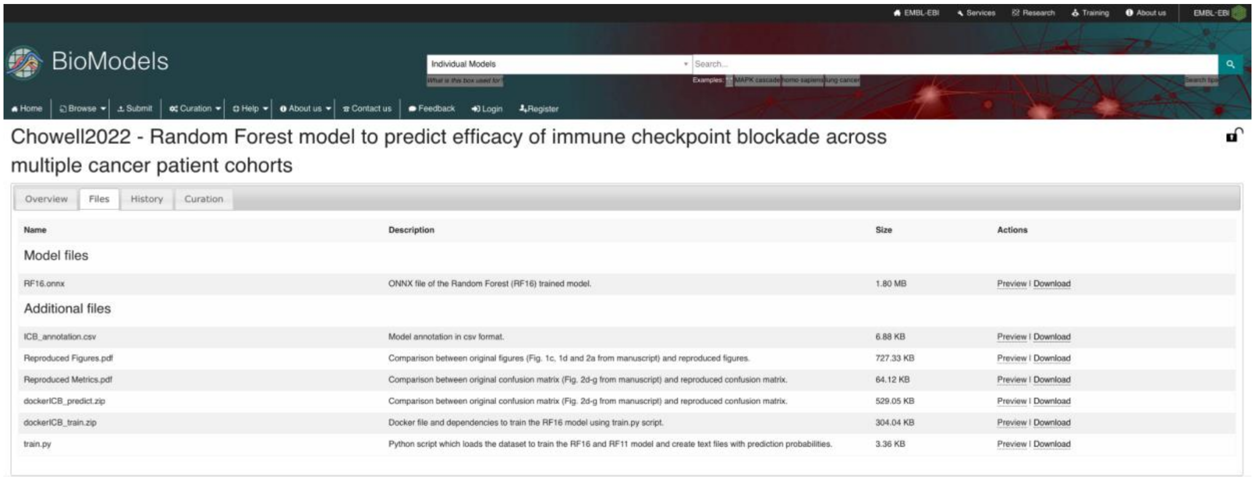


*Supplementary Figure 2: Files Tab of Chowell2022 submitted model.*

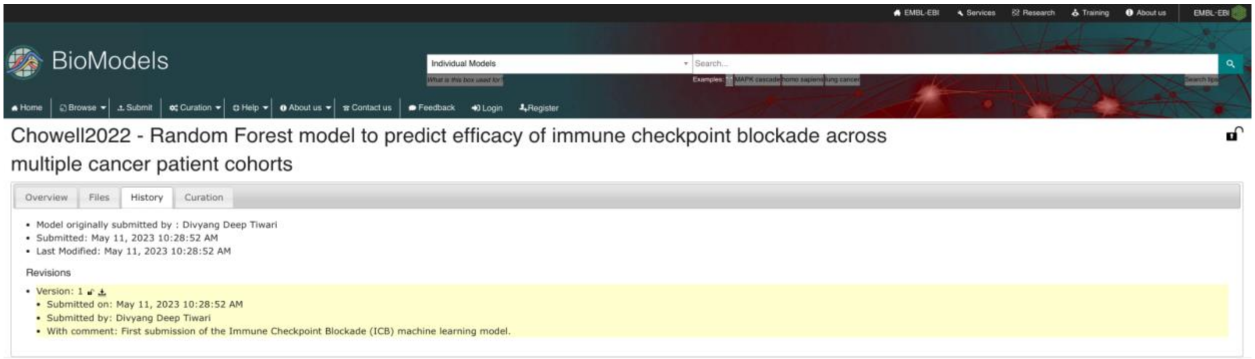


*Supplementary Figure 3: History Tab of Chowell2022 submitted model.*

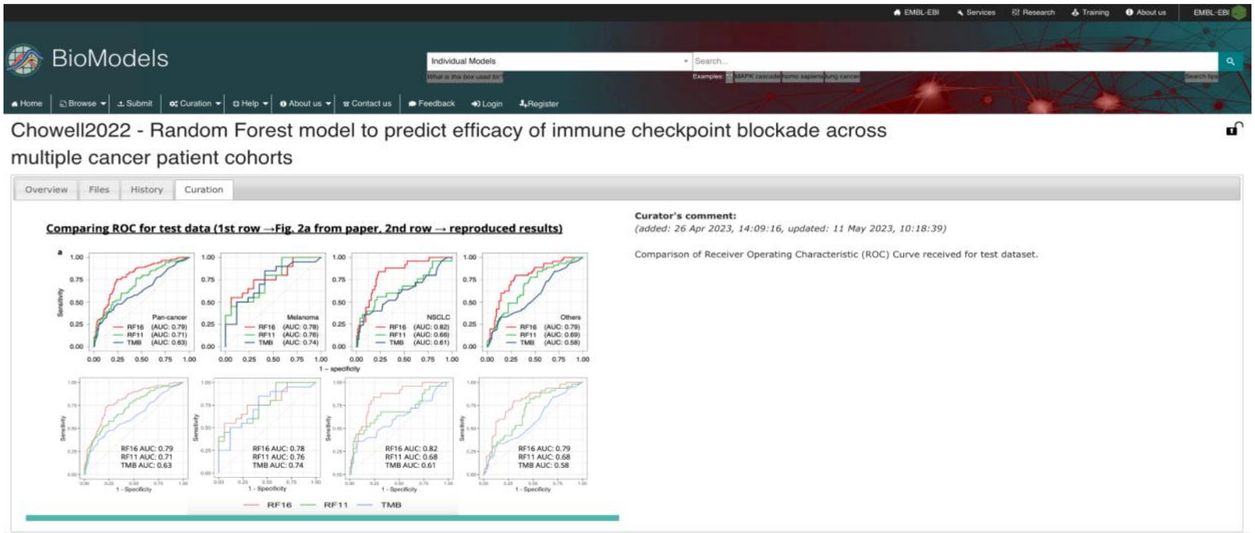


*Supplementary Figure 4: Curation Tab of Chowell2022 submitted model.*

### Use Case #2: Wang et al., 2022, RFDesign for Protein Inpainting and Design

Since many models are too large to fully retrain in an academic setting, but are nevertheless useful for practitioners, we also reproduced such a model, starting from the trained weights and evaluating the model on both the evaluation metrics and examples presented in the publication for reproduction purposes as well as on other related examples. The model in this case was RFDesign (Wang *et al.*, 2022), a fine-tuned version of the structure prediction network aimed to aid with protein design tasks in which parts of the sequence should be redesigned.

After installing the required libraries and downloading the pretrained weights, the tests provided by the authors were run and their tutorial examples successfully reproduced.

A few issues were present at the start of the model reproduction such as missing dependencies and outdated scripts. However, after discussion with the authors these obstacles were removed, enabling the described reproduction. The model files are available at MODEL2304300001.

### Use Case #3: Kong et al., 2023, MEAN for Antibody Design and Affinity Maturation

As a follow-up to the Bio-Hackathon 2022, we wanted to repeat the reproduction process with other models. One such example that was reproduced was the MEAN model by Kong, Huang and Liu (2023), a graph neural network aimed at tasks for antibody design and affinity maturation.

The datasets as well as the training code was obtained from the GitHub repository provided by the authors. The model was successfully trained and the performance reproduced with respect to the three tasks studied by the authors. The data for evaluating the model as well as the evaluation metrics depends on the task:

- Task 1: Generative antibody sequence and structure modelling. For this task, 3,127 complexes from the Structural Antibody Database (SAbDb) were selected, filtered and pre-processed using the provided scripts. Evaluation metrics for this task were amino acid recovery (AAR) and Root Mean Square Deviation (RMSD).
- Task 2: Antigen-binding CDR-H3 Design. The evaluation dataset consists of 60 diverse complexes selected from RAbD. In addition to AAR and RMSD, TM-score was added as a metric.
- Task 3: Affinity optimisation for improvement of existing antibody candidates.

The tasks could be successfully accomplished via the reproduced model. The Dockerfile with all dependencies and scripts needed to reproduce and use the model was submitted to BioModels. However, a conversion of the model to ONNX format was not possible due to the missing ONNX support for several functions inside the neural network architecture (for example logical operations such as torch.logical_and or torch.logical_or). The model files are available at MODEL2304300002.

### Use Case #4: Deshpande 2019, Random Forest model to predict long non-coding RNAs from coding RNAs in Zea Mays plant transcriptomic data

We succeeded in exporting, executing and validating the random forest classifier model of Deshpande *et al.* (2019) during the BioHackathon 2022. The training model consists of 16,000 coding RNA and lncRNA sequences applying a Random Forest classifier using n_estimators of 600 and Gini Impurity as the quality of split criterion. The training and test set sequences were obtained from CANTATAdb v2.0. The Scikit-Learn Python library was used in Python 3.7 to train the model and export the trained ML model in ONNX format. Re-execution of the model using the ONNX-runtime for Intel provided identical results to the native execution using Scikit-Learn, based on a comparison of Precision, Recall, F1-Score and Support (see Supplementary Table 2) on two different datasets P0 and P1.

Additional metadata was added describing the biological context of the model. It was submitted to the BioModels repository as MODEL2304280002.

| | **NM P0** | **ONNX P0** | **NM P1** | **ONNX P1** |
|---|---|---|---|---|
| Precision | 0.95 | 0.95 | 0.9 | 0.9 |
| Recall | 0.90 | 0.90 | 0.95 | 0.95 |
| F1-Score | 0.93 | 0.93 | 0.93 | 0.93 |
| Support | 519 | 519 | 481 | 481 |

*Supplementary Table 2. Comparison of native model (NM) and converted ONNX model (ONNX) on two distinct datasets (P0, P1).*

### Use Case #5: Nassar et al., 2022, Automatic Document Classification & Entity Extraction

In this use case, we attempted to observe whether it follows the FAIR principles in the automatic extraction of entities of bio-literature. Thus, we can find, access, and reuse interoperable data from any such literature. To achieve this, we first replicated the work of Nassar, M. *et al.* (2022). In this work, a machine learning framework enriches various microbiome studies (n=19,209) with essential and accurate metadata from open-access research articles (n=114,099). Classifying the documents to prepare a publication triage for a wide variety of microbiome environments is essential. Nassar *et al.* addressed this requirement by constructing supervised training data sets where the GOLD annotations were assigned to metagenomics studies.

The data sets mapped to the corresponding MGnify (Gurbich, T.A. *et al.*, 2023) cross-referenced publications. The random forest models were trained on the hierarchical levels of GOLD ontology, yielding diverse biome prediction models. The authors used ten documents from Europe PMC (The Europe PMC Consortium, 2015) as a test set. They set a threshold of 0.40 prediction probability for the top GOLD hierarchical levels (Engineered, Environmental, and Host-associated).
In our current task, we replicated the result by annotating and classifying the test documents. Then we converted the classification model into ONNX format. The ONNX model predicted the same

results as the original model (result rounded to two significant digits after decimal). We compared the performance of the original model and the ONNX model. We show the results in the following two graphs (Supplementary Figs. 5 & 6). We show prediction probability for document classification into the Environmental, Host-Associated and Engineered categories in different colours. We show the equivalent results of the models for the 2-digit significance level after the decimal. The model files are available at MODEL2304210001.

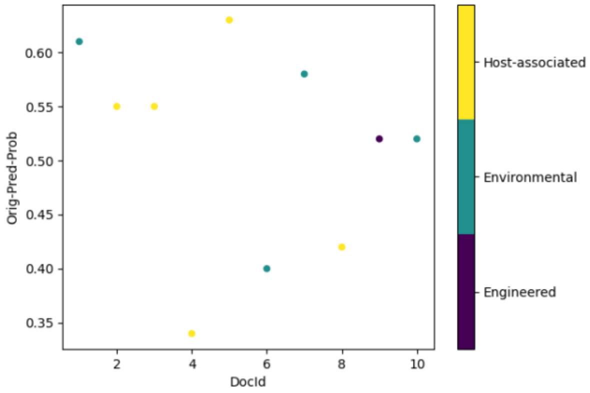


*Supplementary Figure 5: Prediction Probability with original model for 10 documents for three categories (shown in three colours).*

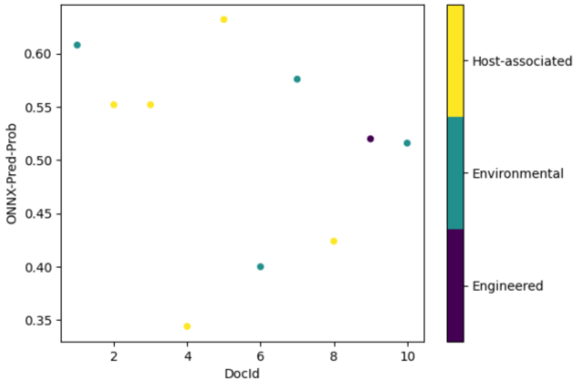


*Supplementary Figure 6: Prediction Probability with ONNX model for 10 documents for three categories (shown in three colours).*

### Use Case #6: Alam and Islam, 2019, Red-Blood Cell Classification using Tiny YOLO v2

During the BioHackathon, we selected a dataset for classification and annotation of red blood cells in microscopy images (Alam and Islam, 2019) based on the (Tiny) YOLO v2 network for object recognition in images. We followed the instructions provided by the authors to install the dependencies and pre-trained model weights and were able to reproduce their first sample image annotation without issues.

After initial attempts to export the pre-trained TensorFlow (TF) 2 model from a checkpoint, we adapted the source code to store the model with weights in the Protocol Buffers serialisation format directly. This could then be read using the Python libraries for TensorFlow to ONNX conversion to generate an equivalent ONNX model. We were then able to create an inference session using the pretrained ONNX model based on the Intel-accelerated ONNX-runtime library which we then applied to one of the test images supplied by the authors of the original model. Some issues were encountered concerning the additional pre- and postprocessing of the red blood cell images, which the authors originally implemented with custom Python code.

We were able to port their code to work with the ONNX model and evaluated the image annotation results on their original test image dataset, against their original ground truth. This resulted in qualitatively identical predictions for the original TF model and the ONNX model (see *Supplementary Table 3*). Some minor differences remained in positions of cell centres and in the sizes of classified regions (see *Supplementary Figures 7 & 8*), but these were minor and may have been caused by slight numerical differences introduced by the ONNX runtime or by rounding differences introduced in our custom code. We further ensured a reproducible codebase by capturing and pinning dependencies using a Conda environment. This was also used as the foundation for the Dockerfiles for training of the original TF model and the application of the trained ONNX model. Model files are available at MODEL2305050001.

*Supplementary Table 3: Total counts of annotated cell types compared between ground truth, TensorFlow 2 and ONNX models. Excess counts for Red Blood Cells (RBCs) for TF2 and ONNX include RBCs that are only partially visible in the microscopy images, while for the ground truth only RBCs that were completely contained within the image boundaries were counted.*

| **Cell Type** | **Ground Truth** | **ONNX** | **TF2** |
|---|---|---|---|
| Platelets | 55 | 54 | 54 |
| RBC | 792 | 853 | 853 |
| WBC | 61 | 60 | 60 |

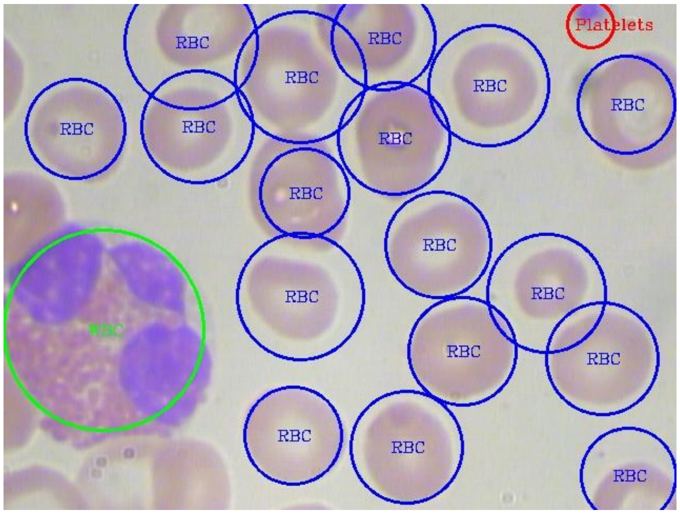


*Supplementary Figure 7: Original blood cell image annotation created using the TF2 implementation.*

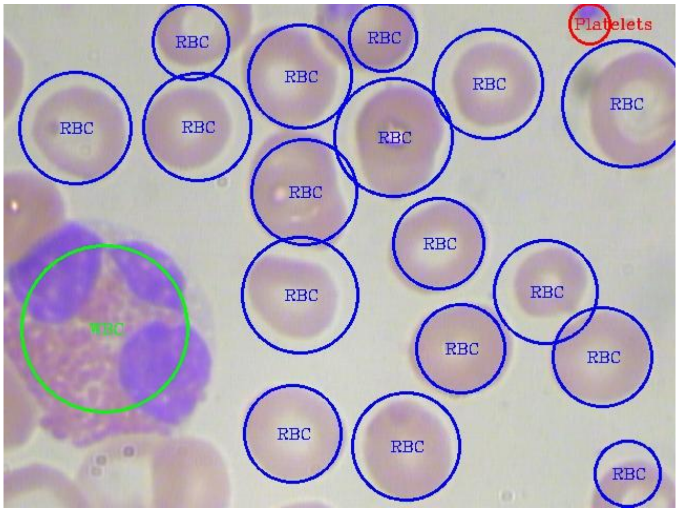


*Supplementary Figure 8: Reproduced blood cell image annotation created using the ONNX model after conversion from the TF2 model. Some differences are visible in the annotation areas, specifically for the white blood cell (WBC) in this example. Annotation area centres are largely identical.*
